# Uterine stromal but not epithelial PTGS2 is critical for murine pregnancy success

**DOI:** 10.1101/2024.10.24.620133

**Authors:** Noura Massri, Ripla Arora

**Author notes:** Corresponding Author: Ripla Arora Associate Professor, Department of Obstetrics, Gynecology and Reproductive Biology Institute for Quantitative Health Science and Engineering Michigan State University, 775 Woodlot Drive, Rm#3312 East Lansing, MI-48824, USA.

## Abstract

Use of non-steroidal anti-inflammatory drugs that target prostaglandin synthase (PTGS) enzymes have been implicated in miscarriage. Further, PTGS2-derived prostaglandins are reduced in the endometrium of patients with a history of implantation failure. However, in the mouse model of pregnancy, peri-implantation PTGS2 function is controversial. Some studies suggest that *Ptgs2^-/-^* mice display deficits in ovulation, fertilization, and implantation, while other studies suggest a role for PTGS2 only in ovulation but not implantation. Further, the uterine cell type responsible for PTGS2 function and role of PTGS2 in regulating implantation chamber formation is not known. To address this we generated tissue-specific deletion models of *Ptgs2*. We observed that PTGS2 ablation from the epithelium alone in *Ltf^cre/+^; Ptgs2^f/f^* mice and in both the epithelium and endothelium of the *Pax2^cre/+^; Ptgs2^f/f^* mice does not affect embryo implantation. Further, deletion of PTGS2 in the ovary, oviduct, and the uterus using *Pgr^cre/+^; Ptgs2^f/f^* does not disrupt pre-implantation events but instead interferes with post-implantation chamber formation, vascular remodeling and decidualization. While all embryos initiate chamber formation, more than half of the embryos fail to transition from blastocyst to epiblast stage, resulting in embryo death and resorbing decidual sites at mid-gestation. Thus, our results suggest no role for uterine epithelial PTGS2 in early pregnancy but instead highlight a role for uterine stromal PTGS2 in modulating post-implantation embryo and implantation chamber growth. Overall, our study provides clarity on the compartment-specific role of PTGS2 and provides a valuable model for further investigating the role of stromal PTGS2 in post-implantation embryo development.

## INTRODUCTION

According to the American College of Obstetricians and Gynecologists, approximately 26% of pregnancies end in miscarriage, and only 10% of these losses are clinically recognized (Bulletins—Gynecology, 2018). Additionally, 1-2% of women experience recurrent pregnancy loss due to undetermined causes (Turesheva et al., 2023). Given the ethical considerations, human pregnancies cannot be studied directly. Thus, mice are often utilized as a model system to understand the early events of pregnancy. Recent advancements in 3D imaging methodology have been successfully applied to the pre-implantation stages of a mouse pregnancy, revealing phenomena that are challenging to uncover using traditional 2D histology. 3D imaging has revealed that embryo clusters enter the uterine environment at gestational day (GD) 3, ∼72 hours after the mouse mating event. These embryos initially move together as clusters towards the middle of the uterine horn and then they undergo a bidirectional scattering movement followed by embryo spacing along the oviductal-cervical axis (Flores et al., 2020, Chen et al., 2013). At GD4, once the embryo arrives in the center of a flat peri-implantation region of the uterine lumen, a V-shaped embryo implantation chamber begins to form (Madhavan et al., 2022). This is concurrent with increased vascular permeability and sprouting angiogenesis at the embryo implantation sites at GD4 1800h (Madhavan et al., 2022, Massri et al., 2023). The proper formation of the embryo implantation chamber is critical as it facilitates embryo alignment along the mesometrial-anti-mesometrial axis, where the blastocyst’s inner-cell mass faces the uterine mesometrial pole (Madhavan et al., 2022). Following embryo implantation, decidualization occurs, where stromal cells in the uterus become epithelialized, and embryos grow to the epiblast stage at GD5. Aberrations in events surrounding embryo implantation and decidualization can lead to a cascade of events that negatively impact subsequent pregnancy development, ultimately resulting in miscarriage and pregnancy loss (Cha et al., 2012).

Successful embryo implantation and maintenance of early pregnancy rely on a delicate interplay of numerous molecular mechanisms (Chen et al., 2013). Among these, prostaglandins (PGs), PGE_2_, PGI_2_, and PGF_2_ have emerged as critical mediators of reproductive success (Wang and Dey, 2006, Psychoyos et al., 1995, Clark and Myatt, 2008). PG synthesis begins with the phospholipase A2 enzyme cleaving arachidonic acid from the phospholipid bilayer. The prostaglandin synthase enzyme 1 (PTGS1) and PTGS2 convert arachidonic acid to PGH2. PGH2 is then converted to PGD_2_, PGE_2_, PGF_2_α, PGI_2,_ and thromboxane by PGD synthase (Funk, 2001). Both PTGS1 and PTGS2 are glycosylated proteins with two catalytic sites: peroxidase and cyclooxygenase (thus the alternate names COX1 and COX2). These enzymes are similar at the amino acid level, but PTGS2 has an extra “side pocket” that allows more space in the active site for substrate binding (Vecchio and Malkowski, 2011). PTGS2 is often induced by cytokines, growth factors, hormones, inflammation, and embryo attachment (Chakraborty et al., 1996, Ricciotti and FitzGerald, 2011), while PTGS1 is constitutively expressed (Ricciotti and FitzGerald, 2011).

Numerous studies have found evidence of PTGS1 and PTGS2 expression in human uterine compartments during implantation (Marions and Danielsson, 1999). PTGS1 is expressed at a constant level in the human endometrium, while PTGS2 is expressed explicitly in the glandular epithelial cells and the endothelial cells (Marions and Danielsson, 1999), and the stromal cells (Stavreus-Evers et al., 2005). Additionally, there is evidence for both PTGS1 and PTGS2 expression in the uteri of various species, including mice (Chakraborty et al., 1996), western spotted skunks (Das et al., 1999), baboons (Kim et al., 1999), and hamsters (Evans and Kennedy, 1978, Wang et al., 2004b). PTGS2 is expressed in the luminal epithelium and sub-epithelial stroma surrounding the blastocyst attachment site in the anti-mesometrial pole, and its expression is induced by the presence of the embryo (Chakraborty et al., 1996). Post embryo implantation, PTGS1 is expressed in the secondary decidual zone; however, PTGS2 expression is localized at the mesometrial pole (Chakraborty et al., 1996).

Non-steroidal anti-inflammatory drugs (NSAIDs) that block PTGS1 and PTGS2 function are amongst the most common over-the-counter medications that women take during pregnancy (Thorpe et al., 2013). There is evidence for an 80% increased risk of miscarriage with the consumption of NSAIDs during pregnancy (Li et al., 2003, Li et al., 2018a, Jackson-Northey and Evans, 2002). PTGS1 has not been shown to have a role in pregnancy in women, and PTGS1-deficient mice do not display significant reproductive issues during pregnancy, except for prolonged parturition (Langenbach et al., 1995). On the other hand, studies in pregnant women who experience recurrent pregnancy loss or implantation failure after in-vitro fertilization procedures demonstrate dysregulation in endometrial PTGS2 (Achache et al., 2010), and its derived prostaglandin PGI2 (Wang et al., 2010). Furthermore, genetic variations in the PTGS2 gene are associated with an increased risk of implantation failure among women going through assisted reproductive procedures (Salazar et al., 2010). In rodents, Lim et. al determined that PTGS2-deficient mice are infertile due to ovulation, fertilization, and implantation deficits (Lim et al., 1997). While ovulation and fertilization defects are widely accepted, there is a controversy regarding the role of PTGS2 during embryo implantation (Lim et al., 1997, Cheng and Stewart, 2003). Chang et. al reported that when wild-type blastocysts are transferred into PTGS2-deficient pseudo pregnant uteri, a 24-hour delay in decidualization is observed, but pregnancy proceeds to birth normally (Cheng and Stewart, 2003). These data suggest that PTGS2 may not be essential for implantation, decidualization, and overall pregnancy success. To explain the discrepancy between these studies it has been proposed that mixed mouse genetic background allows the upregulation of PTGS1 in PTGS2-deficient animals and this PTGS1 may compensate for the loss of PTGS2 (Wang et al., 2004a).

To resolve the controversy surrounding the function of PTGS2 in embryo implantation and to determine the compartment in which PTGS2 function is essential, we utilized the cre-lox recombinase system (Kim et al., 2018). We deleted PTGS2 in the adult uterine epithelium using *Ltf^cre/+^*(Daikoku et al., 2014), in the embryonic uterine epithelium and endothelium using *Pax2^cre/+^(Ohyama and Groves, 2004, Granger et al., 2023)*, and in the epithelial and stromal compartment of the uterus using *Pgr^cre/+^*(Soyal et al., 2005, Madhavan and Arora, 2022) (**Table 1**). We determine that PTGS2 function in the uterine epithelium and endothelium is not critical for implantation or pregnancy success. However, stromal PTGS2 is critical for post-implantation embryo and implantation chamber growth for continued pregnancy progression.

**Table 1:** Mouse models used to study PTGS2 function in the uterus during implantation. . The table outlines the models utilized in the study, along with the corresponding tissues and times of deletion for PTGS2. *Ltf^cre/+^; Ptgs2^f/f^* deletes PTGS2 in the uterine luminal and glandular epithelium in adult mice > 10 weeks. *Pax2^cre/+^; Ptgs2^f/f^*deletes PTGS2 in the uterine luminal, glandular epithelium, and uterine endothelium at the embryonic stage. *Pgr^cre/+^; Ptgs2^f/f^* deletes PTGS2 in the ovary, oviduct, and uterine epithelium (luminal and glandular epithelium), circular smooth muscle, and stroma during neonatal stages. LE: Luminal Epithelium, GE: Glandular Epithelium.

| Model | Ovarian/Oviduct/Uterine Compartment | Deletion time |
| --- | --- | --- |
| *Ltf<sup>cre/+</sup>; Ptgs2<sup>f/f</sup>* | Uterine epithelium (LE, GE) | Adult (> 8.5 weeks) (Daikoku et al, 2014) |
| *Pax2<sup>cre/+</sup>; Ptgs2<sup>f/f</sup>* | Uterine epithelium (LE, GE) and endothelium | Embryonic (GD11.5) (Ohyama & Groves, 2004) |
| *Pgr<sup>cre/+</sup>; Ptgs2<sup>f/f</sup>* | Ovarian granulosa cells, oviductal epithelium and myometrium, uterine epithelium (LE, GE), circular smooth muscle and stroma | Neonatal (P5 epithelium, stroma, P21 circular muscle)(Madhavan & Arora, 2022; Soyal et al, 2005) |

## RESULTS

### Peri-implantation PTGS2 expression in embryo mediated and in oil-stimulated pregnancy

To determine which uterine cells might contribute to PTGS2 expression during peri-implantation stages we performed expression analysis of PTGS2 in the uterine tract during peri-implantation stages utilizing natural and artificial pregnancy models. At GD3 1600h, when embryos are present in the uterus, PTGS2 is not expressed in the uterine luminal epithelium (**Supplementary Fig. 1A, A’**). mRNA expression of *Ptgs2* has been reported in the luminal epithelium when a pseudopregnant uterus is stimulated with oil (Lim et al., 1997). We also observed PTGS2 protein expression in the uterine luminal epithelium four hours after intraluminal oil stimulation of the pseudopregnant uterus at GD3 1200h (**Supplementary Fig. 1B, B’**). At GD4 1200h, when the embryo is at the center of the peri-implantation region, PTGS2 is expressed only in the luminal epithelium but not in the stroma (Madhavan et al., 2022). Following embryo implantation at GD4 1800h, PTGS2 is observed in the uterine sup-epithelial stroma surrounding the embryo implantation chamber (**Supplementary Fig. 1C, C’**), as reported previously (Chakraborty et al., 1996, Madhavan et al., 2022). At GD5.5, PTGS2 is expressed at the mesometrial pole surrounding the embryo implantation chamber as reported previously (Chakraborty et al., 1996) and uterine glands at the implantation chamber (**Supplementary Fig. 1D, D’**).

### PTGS2 deletion in the uterine luminal epithelium and endothelium does not affect embryo implantation, embryo growth, and pregnancy progression

To determine if the uterine epithelium is responsible for pre-implantation PTGS2 function, we generated tissue-specific deletion models of PTGS2 using cre-lox recombinase methodology (Kim et al., 2018) (**Supplementary Fig. 2A, B**). For adult uterine epithelial deletion, we used *Ltf^cre/+^ ; Ptgs2^f/f^* mice (Daikoku et al., 2014) (**Table 1**), and for embryonic uterine epithelium and endothelial deletion, we used *Pax2^cre/+^; Ptgs2^f/f^* mice (Ohyama and Groves, 2004) (**Table 1**). To confirm PTGS2 depletion in the CDH1 positive uterine epithelial cells we used oil-stimulated pseudopregnancies for both *Ltf^cre/+^; Ptgs2^f/f^*, and *Pax2^cre/+^; Ptgs2^f/f^* models (**Fig. 1A, A’, B, B’, C, C’)**. At GD4 1800h, we observed the formation of the V-shaped embryo implantation chamber and stromal PTGS2 expression in control, *Ltf^cre/+^; Ptgs2^f/f^*, and *Pax2^cre/+^; Ptgs2^f/f^* mice (**Fig. 1D, D’, E, E’, F, F’**). At GD4 1800h, we observed no defects in the development of the blastocyst in the *Pax2^cre/+^; Ptgs2^f/f^* uteri **(Fig. 3G, H, and Table 3**). Epithelial-specific and epithelial and endothelial-specific PTGS2-deficient mutants displayed normal embryo spacing and increased vessel permeability at embryo implantation sites, as observed by the blue dye reaction at GD4 (**Fig. 1I, JK).** At GD12.5 we observed that uteri from both mutants displayed embryos that had developed similar to embryos from control uteri (**Fig. 1J, L**). Further, both *Ltf^cre/+^; Ptgs2^f/f^* and *Pax2^cre/+^; Ptgs2^f/f^*mice were able to go to term with no significant effect on the duration of the pregnancy or the number of the pups born (**Fig. 1K, L**). Overall, our data suggest that the uterine epithelium and endothelium are not the sources of PTGS2-derived prostaglandin synthesis critical for implantation and pregnancy progression.

**Figure 1.**
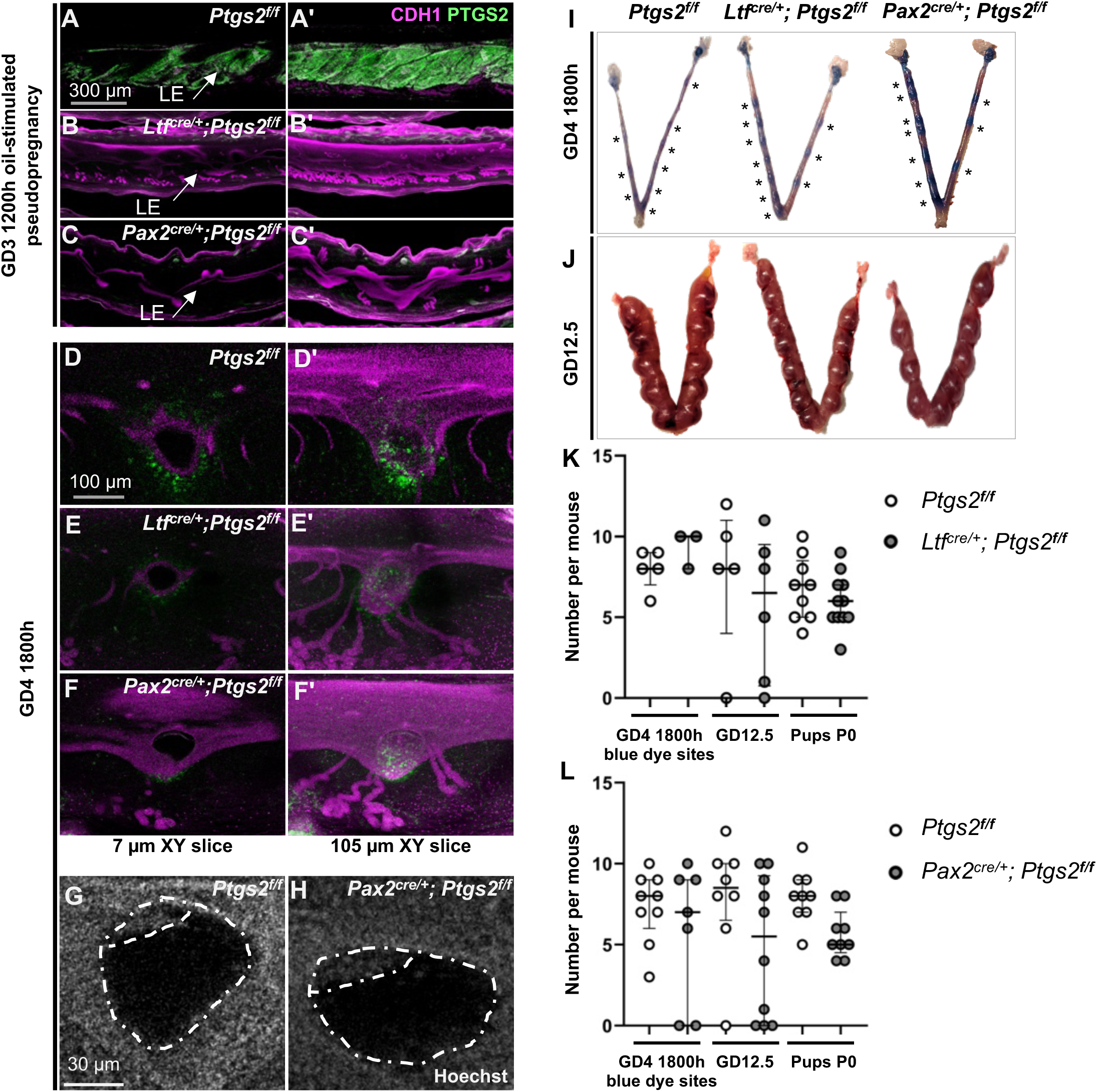
Conditional deletion of PTGS2 in the uterine epithelium and endothelium does not affect embryo implantation and pregnancy success. PTGS2 expression in CDH1 positive cells in oil-stimulated pseudo pregnant *Ptgs2^f/f^* uteri (A), *Ltf^cre/+^; Ptgs2^f/f^* uteri (B) and *Pax2^cre/+^; Ptgs2^f/f^* uteri (C) at GD3 1200h, 4h after intraluminal oil stimulation. 3 different regions from at least 4 uterine horns were evaluated. 7µm XY slice (A, B, C). 105µm XY slice (A’, B’, C’). PTGS2 expression in the subepithelial stroma in *Ptgs2^f/f^* (D), *Ltf^cre/+^; Ptgs2^f/f^* (E), and *Pax2^cre/+^; Ptgs2^f/f^* uteri (F) at GD4 1800h. 7µm XY slice (D, E, F). 105µm XY slice (D’, E’, F’). At least 2 implantation sites from 3 different uterine horns were analyzed. The top of the images represents the mesometrial pole, while the bottom represents the anti-mesometrial pole. Blastocyst stage embryos in *Ptgs2^f/f^* (G) and *Pax2^cre/+^; Ptgs2^f/f^* mice (H) at GD4 1800h. White dashed lines: blastocyst. Uteri with blue dye sites at GD4 1800h (I). Black asterisks: bluedye sites. Uteri with embryo sites at GD12.5 (J). Quantitation of blue dye sites at GD4 1800h, live embryos at GD12.5, and P0 pups in *Ltf^cre/+^; Ptgs2^f/f^* mice (K), and in *Pax2^cre/+^; Ptgs2^f/f^*(L) with their respective controls. Each dot represents one mouse analyzed. Median values shown. Data analyzed using unpaired parametric t-test. No significant differences were observed. Scale bars, A-C’: 300 µm, D-F’: 100 µm, G, H: 30 µm. LE: Luminal epithelium.

### Stromal deletion of PTGS2 results in mid-gestation decidual resorption

To delete *Ptgs2* in the granulosa cells of the pre-ovulatory follicle and the corpus luteum, the epithelium, and the myometrium of the oviduct (Soyal et al., 2005), and the circular smooth muscle, epithelium, and stroma of the uterus (Soyal et al., 2005, Madhavan and Arora, 2022) we utilized the Progesterone-Receptor-driven Cre (*Pgr^cre^)* mouse line (**Table 1, and Fig. 2A, A’, B, B’**). We observed normal embryo spacing in *Pgr^cre/+^; Ptgs2^f/f^* mice; however, embryo implantation was delayed as observed using the blue dye reaction at GD4 1800h (**Fig. 2C, D, G**) (median blue dye sites in controls: 10, *Pgr^cre/+^; Ptgs2^f/f^*: 7, P<0.05). 24 hours later at GD5.5, a similar number of decidual sites was observed in controls and *Pgr^cre/+^; Ptgs2^f/f^* uteri (**Fig. 2D, G**). To determine the cause for delayed implantation in the mutant mice, we determined the mRNA expression of a critical glandular cytokine, *Leukemia inhibitory factor* (*Lif*) at GD3 1800h. We observed reduced levels of *Lif* mRNA in FOXA2+ glandular epithelial uterine cells in *Pgr^cre/+^; Ptgs2^f/f^* uteri (**Supplementary Fig. 3A, B, C**). However, we observed no differences in the serum progesterone levels between control and mutant mice at GD3 and GD4 1800h (**Supplementary Fig. 3D**). Similar to GD5.5, at GD8.5, we observed no significant difference in the number of decidual sites between control and *Pgr^cre/+^; Ptgs2^f/f^* uteri; however, we started to observe a few resorption sites in the mutants **(Fig. 2E, H**). At GD12.5, the number of decidual sites was similar; however, we observed a significant number of resorbing decidua (50%) in the mutant uteri (median live embryo number in control: 9, *Pgr^cre/+^; Ptgs2^f/f^*: 4, P<0.01) (**Fig. 2F, H**). Commensurate with the resorptions at mid-gestation, we observed a significant reduction in pups born to *Pgr^cre/+^; Ptgs2^f/f^*females (46% pups loss) in comparison with control (median live pup number in controls: 8, *Pgr^cre/+^; Ptgs2^f/f^*: 4, P<0.05) (**Fig. 2I**).

**Figure 2.**
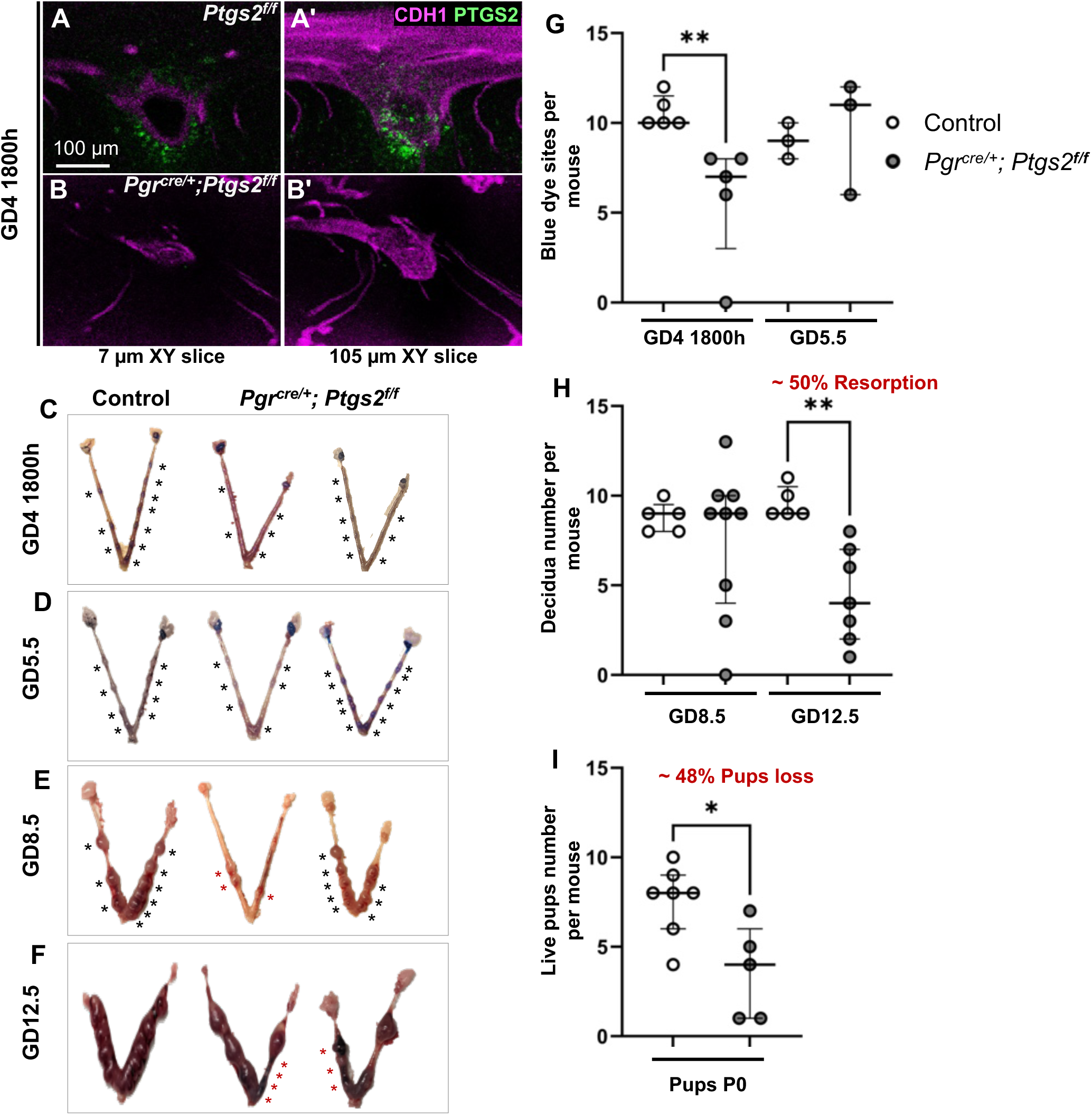
*Pgr^cre/+^; Ptgs2^f/f^* mice display a delay in embryo implantation, mid-gestation decidual resorption, and pregnancy loss. PTGS2 expression in the subepithelial stroma surrounding the embryo implantation chamber in *Ptgs2^f/f^* (A) and *Pgr^cre/+^; Ptgs2^f/f^* (B) uteri at GD4 1800h. At least 9 implantation sites were evaluated in at least 2 mice. 7µm XY slice (A, B). 105µm XY slice (A’, B’). The top of the images represents the mesometrial pole, while the bottom represents the anti-mesometrial pole. Blue dye sites at GD4 1800h (C) and GD5.5 (D). Decidual sites at GD8.5 (E), and GD12.5 (F) in control and *Pgr^cre/+^; Ptgs2^f/f^* uteri. Black asterisks: blue dye sites. Orange arrowheads: resorbed deciduae sites. Quantification of blue dye sites at GD4 1800h and GD5.5 (G), decidual sites number at GD8.5 and at GD12.5 (H) and live pups at P0 (I) in both groups. At least n=3 mice were evaluated per genotype for each pregnancy stage. Each dot represents one mouse. Median values shown. Data analyzed using unpaired parametric t-test. * P < 0.05, ** P < 0.01. Scale bar for A-B’: 100 µm.

### Abnormal embryo development in the post-implantation chamber of PTGS2-deficient uteri

To determine the first time point when embryo development is affected in the *Pgr^cre/+^; Ptgs2^f/f^* uteri, we examined embryo morphology at different time points during gestation. *Ptgs2^-/-^* mice display a ∼30% reduction in the number of eggs ovulated per mouse and a complete failure of fertilization (Lim et al., 1997) Thus, we first examined the fraction of fertilized eggs in the *Pgr^cre/+^; Ptgs2^f/f^* mice. We performed an oviductal flush at GD1 1200h and cultured the embryos in vitro for 72 hours. In control mice, we observed that 97.5% of the embryos were at the 2-cell stage at the time of the oviductal flush. After 72 hours of in-vitro embryo culture, 18/39 (45%) embryos reached the morula stage, and 21/39 (52.5%) reached the blastocyst stage (**Table 2 and Fig. 3A, C, E, G, I, J)**. With *Pgr^cre/+^; Ptgs2^f/f^* mice, we observed 12/56 (21.42%) unfertilized eggs, 4/56 (7.14%) 1-cell stage embryos, and 40/56 (71.42%) 2-cell stage embryos at the time of oviductal flush. After 72 hours of in-vitro embryo culture, 8/56 (14.28%) embryos reached the morula stage, and 36/56 (64.28%) embryos reached the blastocyst stage. The 12/56 (21.42%) unfertilized eggs remained as such with no extrusion of polar body and cell division (**Table 2 and Fig. 3B, D, F, H, I, J**). Thus, *Pgr^cre/+^; Ptgs2^f/f^* mice show a substantially improved fertilization rate compared to *Ptgs2^-/-^* mice (Lim et al., 1997, Matsumoto et al., 2001). Overall, in our *Pgr^cre/+^; Ptgs2^f/f^* model, we noted that once fertilization occurs, these embryos develop normally to the morula/blastocyst stage in vitro.

**Figure 3.**
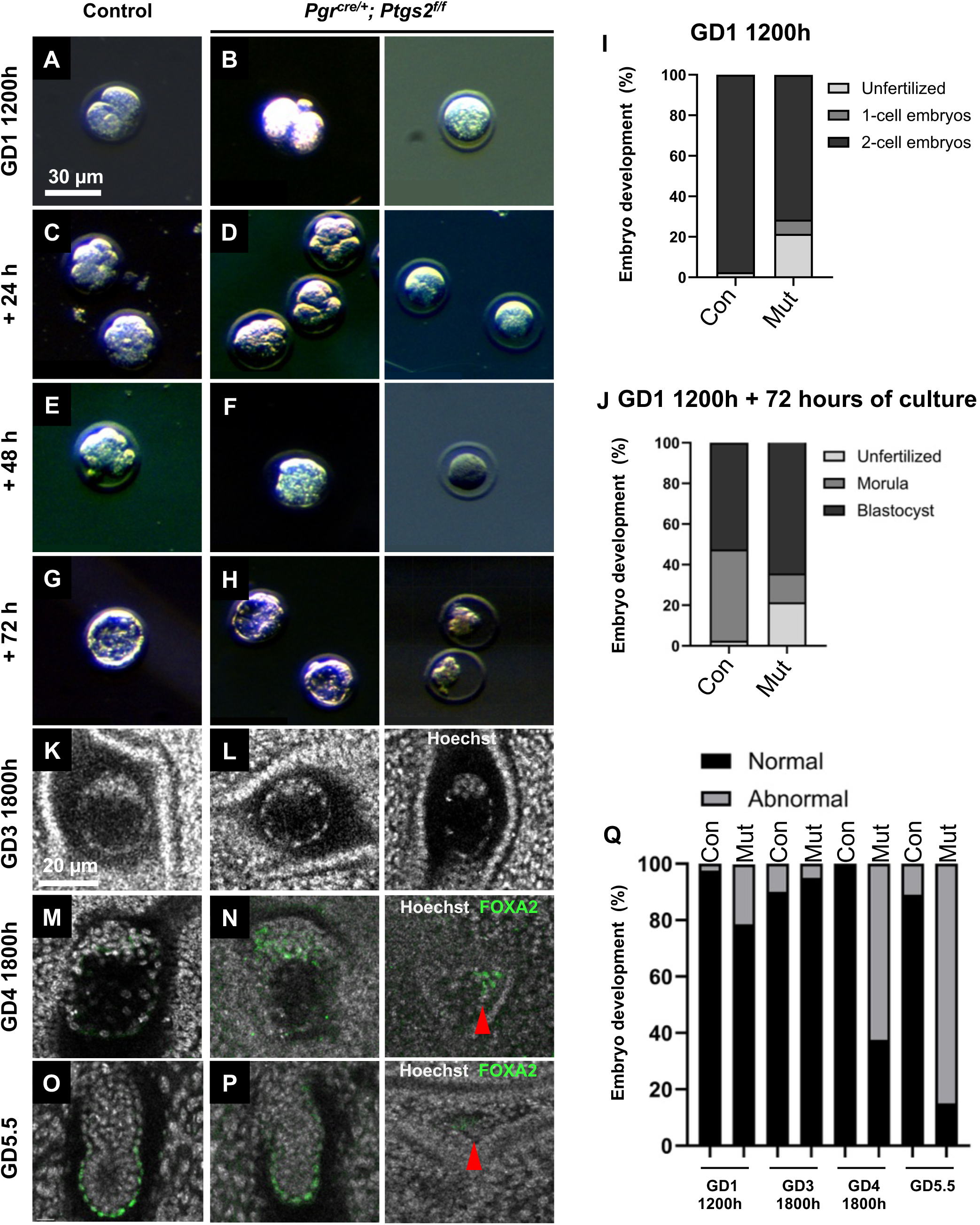
Stromal ablation of PTGS2 restricts embryo growth at post-implantation stages. Oviductal flush at GD1 1200h revealed 2-cell stage embryos in control (A), and 2-cell stage embryos and unfertilized eggs in *Pgr^cre/+^; Ptgs2^f/f^* mice (B). 24, 48, and 72 hours culture of flushed embryos/eggs in control (C, E, G) and *Pgr^cre/+^; Ptgs2^f/f^* mice (D, F, H). Embryo development percentage at GD1 1200h (I) and at GD1 1200h + 72 hours of culture (J). Blastocyst stage embryos in control (K) and *Pgr^cre/+^; Ptgs2^f/f^* mice (L) at GD3 1800h. Blastocyst stage embryos in control (M), and blastocyst and abnormal embryos in *Pgr^cre/+^; Ptgs2^f/f^* mice at GD4 1800h (N). Epiblast stage embryos in control mice (O); and epiblast and abnormal embryos in *Pgr^cre/+^; Ptgs2^f/f^* mice at GD5.5 (P). Red arrowheads: resorbing embryos. Comparison of embryo development percentage across GD1.5 - GD5.5 (Q). Analysis was performed in uteri with embryos. At least n=3 mice were analyzed per time point. Scale bars, A-H: 30 µm, K-P: 20 µm. Con: Control; Mut: mutant *Pgr^cre/+^; Ptgs2^f/f^*.

**Table 2:** In-vitro-embryo culture of embryos flushed from *Ptgs2^f/f^* and *Pgr^cre/+^; Ptgs2^f/f^* uteri at GD1 1200h.

|  | Day of collection/ oviduct flush |  |  | After 72 hours of culture |  |  |
| --- | --- | --- | --- | --- | --- | --- |
| Genotype/<br>Mice number | Unfertilized<br>eggs<br>n (%) | 1-Cell<br>embryos<br>n (%) | 2-cell<br>embryos<br>n (%) | Unfertilized<br>eggs<br>n (%) | Morula<br>n (%) | Blastocyst<br>n (%) |
| *Ptgs2<sup>f/f</sup>*<br>n= 5 | 1 (2.5%) | 0 (0%) | 39 (97.5%) | 1 (2.5%) | 18 (45%) | 21 (52.5%) |
| *Pgr<sup>cre/+</sup>;*<br>*Ptgs2<sup>f/f</sup>*<br>n=8 | 12 (21.43%) | 4 (7.14%) | 40 (71.43%) | 12 (21.43%) | 8 (14.28%) | 36 (64.28%) |

Next, we evaluated embryo development in our *Pgr^cre/+^; Ptgs2^f/f^* model in vivo. We observed that for uteri with embryos, at GD3 1800h, 95% of the embryos reached the blastocyst stage (**Table 3 and Fig. 3K, L**). However, at post-implantation stages at GD4 1800h, we observed that ∼62.5% of the embryos displayed embryo morphology that deviated from the typical elongated blastocyst (**Table 3 and Fig. 3M, N**). At GD5.5, we observed that 85% of the decidual sites had degrading embryos suggestive of pregnancy arrest (**Table 3 and Fig. 3O-P**). Our data suggests that embryonic growth restriction begins soon after implantation in *Pgr^cre/+^; Ptgs2^f/f^* mice (**Table 3 and Fig. 3 Q**).

**Table 3:** Embryo development at GD3 1800h, GD4 1800h, and GD5 1800h in *Ptgs2^f/f^* and *Pgr^cre/+^; Ptgs2^f/f^* mice.

| Stage | Genotype | Mice Number | Uterine horns | Avg. Embryo sites (observed by blue dye) | Embryo sites (Examined by 3D) | Blastocyst or Epiblast<br>n (%) | Abnormal<br>n (%) |
| --- | --- | --- | --- | --- | --- | --- | --- |
| GD3 | *Ptgs2<sup>ff</sup>* | N = 5 | 5 | NA | 20 | 18 (90) | 2 (10) |
|  | *Pgr<sup>cre/+</sup>;*<br>*Ptgs2<sup>ff</sup>* | N = 4 | 5 | NA | 19 | 18 (95) | 1 (5) |
| GD4 | *Ptgs2<sup>ff</sup>* | N = 4 | 4 | 10.6 | 14 | 14 (100) | 0 (0) |
|  | *Pgr<sup>cre/+</sup>;*<br>*Ptgs2<sup>ff</sup>* | N = 5 | 5 | 5.8 | 24 | 9 (37.5) | 15 (62.5) |
|  | *Pax2<sup>cre/+</sup>;*<br>*Ptgs2<sup>ff</sup>* | N = 4 | 5 | 5.85 | 32 | 29 (90.6) | 3 (9.4) |
| GD5 | *Ptgs2<sup>ff</sup>* | N = 3 | 3 | 9 | 9 | 8 (88.88) | 1 (11.11) |
|  | *Pgr<sup>cre/+</sup>;*<br>*Ptgs2<sup>ff</sup>* | N = 5 | 5 | 9.6 | 20 | 3 (15) | 17 (85) |

### Loss of stromal PTGS2 results in an abnormal implantation chamber, reduced implantation site vascular remodeling, and a poor decidualization response

Since we observed defects in the post-implantation embryo, we hypothesized that implantation chamber and decidualization were the critical processes affected by the loss of PTGS2. We reconstructed the implantation chamber at GD4.5 and GD5.5 using 3D confocal imaging and image segmentation. At GD4 1800h, 13/14 embryos in control mice displayed a V-shaped chamber; however, in *Pgr^cre/+^; Ptgs2^f/f^* mice, only 6/24 implantation chamber displayed a V-shape while the remaining 18/24 embryos displayed either an asymmetric or an abnormal V-shaped chamber (**Fig. 4A, B, C)**. At GD5.5, control mice displayed continued elongation of the V-shape chamber while the chambers in the *Pgr^cre/+^; Ptgs2^f/f^* uteri appeared shorter (**Fig. 4D, E, F**). The length of the implantation chamber in *Pgr^cre/+^; Ptgs2^f/f^* mice was significantly lower than control at both GD4 1800h (median chamber length in controls: 575.5µm, *Pgrcre/+; Ptgs2f/f*: 425.5µm, P < 0.001) and GD5.5 (median chamber length in controls: 1007µm, *Pgrcre/+; Ptgs2f/f*: 589.5µm, P < 0.0001) (**Fig. 4G**).

**Figure 4:**
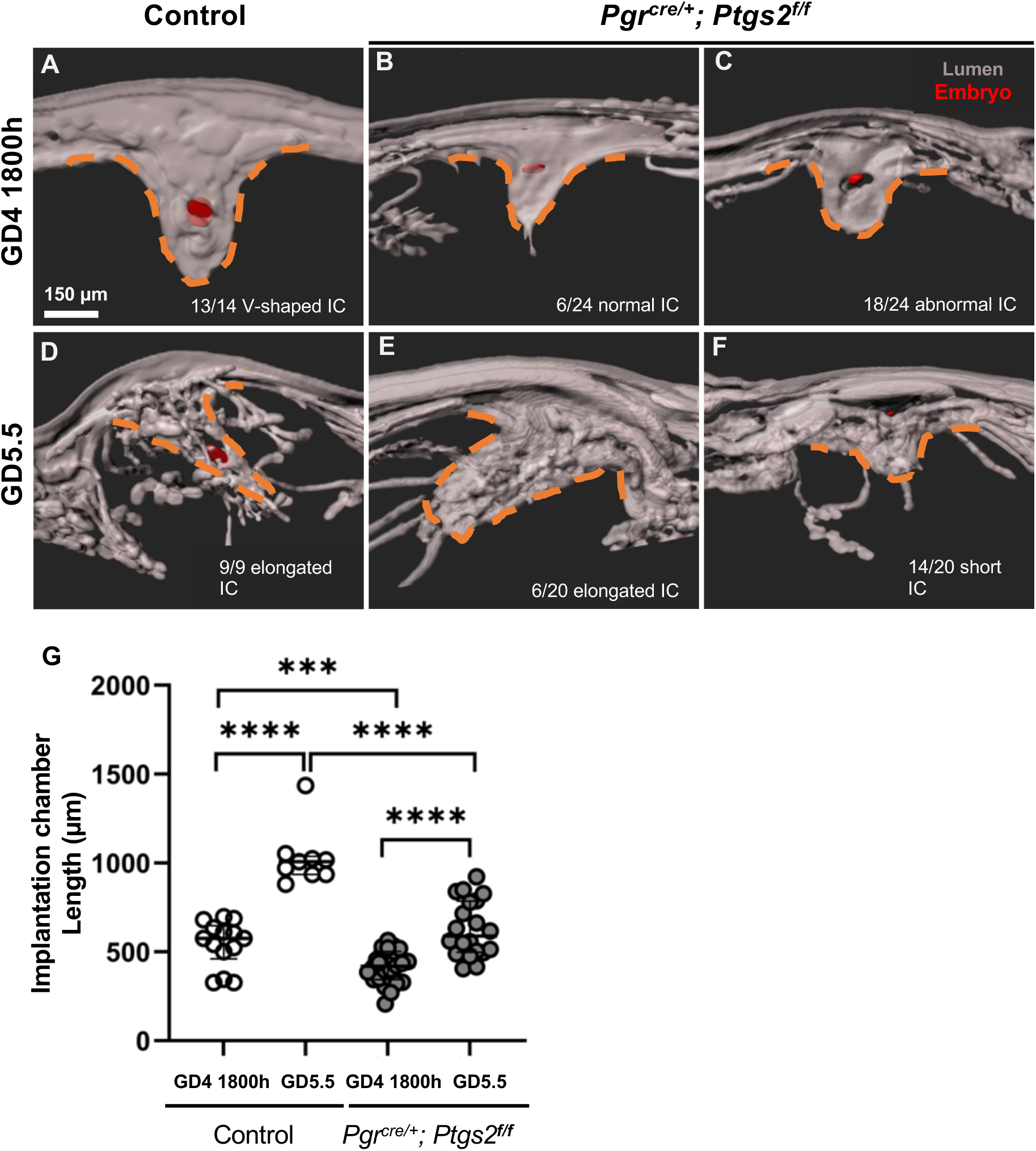
Abnormal embryo implantation chamber structure in *Pgr^cre/+^; Ptgs2^f/f^* mice. At GD4 1800h, V-shaped implantation chambers (13/14) are observed in control mice (A) and 6/24 normal V-shaped implantation chambers (B) and 18/24 abnormally shaped implantation chambers (C) are observed in *Pgr^cre/+^; Ptgs2^f/f^*mice. At GD5.5, elongated embryo implantation chambers (9/9) are observed in control mice (D) and 6/20 elongated (E) and 14/20 short implantation chambers (F) are observed in *Pgr^cre/+^; Ptgs2^f/f^* mice. The top of the images represent the mesometrial pole while the bottom represent the anti-mesometrial pole. Quantitation of implantation chamber in control and *Pgr^cre/+^; Ptgs2^f/f^* mice at GD4 1800h and GD5.5 (G). At least n=3 mice were evaluated per time point. Each dot represents one implantation chamber. Median values shown. Data analyzed using unpaired parametric t-test. *** P < 0.001, **** P < 0.0001. Scale bar, A-F: 150 µm. Orange dashed lines: embryo implantation chamber; IC: Implantation Chamber.

We also evaluated vascular development in the implantation and inter-implantation regions of the uterine horn at GD4 1800h. We observed a drastic decrease in vessel density surrounding the embryo implantation chamber in the mutant uteri compared to controls, however, the vessel density in the inter-implantation site remained comparable (**Fig. 5A, B, C, D**). Vessel diameter was similar in controls and mutants across both implantation and inter-implantation sites (**Fig. 5A, B, D**). CD31-positive cells accumulate around the implantation chamber (Govindasamy et al., 2021), and this expression overlaps with the PTGS2 expression domain. We observed that 8/8 implantation sites in control mice showed this CD31 signal around the implantation chamber (**Fig. 5E, E’, H**), while only 4/10 implantation sites in the mutant showed a CD31 signal around the chamber (**Fig. 5F, F’, G, G’, H**).

**Figure 5:**
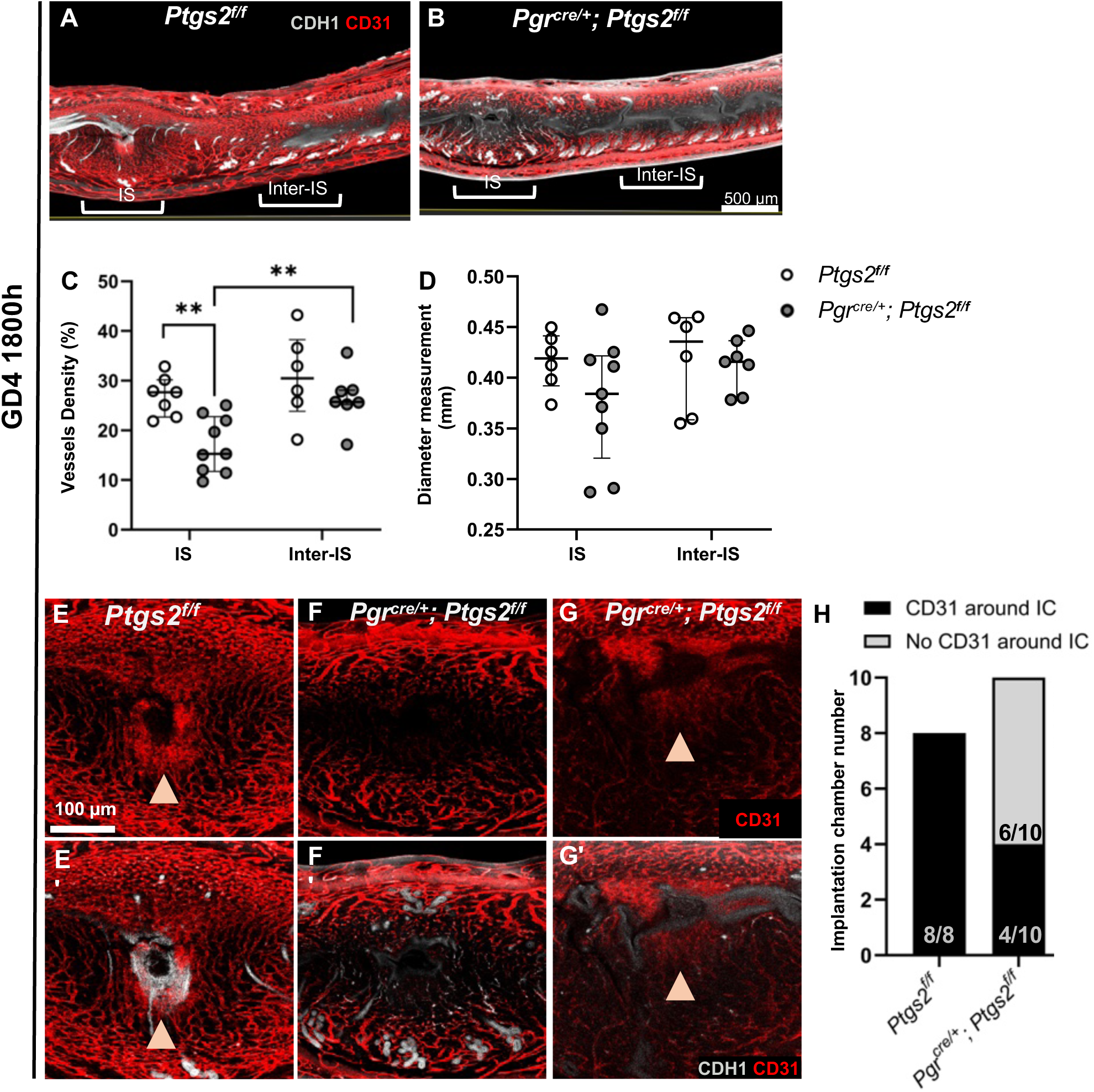
Abnormal vascular development at implantation site in *Pgr^cre/+^; Ptgs2^f/f^.* CD31 expression in *Ptgs2^f/f^* (A) and *Pgr^cre/+^; Ptgs2^f/f^* (B) mice at GD4 1800h. Quantitation of vessel density (C) and vessel diameter (D) at embryo implantation sites and in inter-implantation sites (region between two implantation sites). CD31 expression around the embryo implantation chamber in *Ptgs2^f/f^* (E, E’) and *Pgr^cre/+^; Ptgs2^f/f^* mice (F, F’, G, G’). The top of the images represent the mesometrial pole while the bottom represent the anti-mesometrial pole. Quantification of embryo implantation chamber with and without CD31 expression (H). n=3 mice were evaluated per genotype. Each dot represents one implantation or inter implantation site. Median values shown. Data analyzed using unpaired parametric t-test. ** P < 0.01. Scale bar, A-B: 200 µm, E-G’: 100 µm. IS: Implantation site; Inter-IS: inter-implantation site; IC: implantation chamber.

Given the defects in implantation chamber, we evaluated the expression of classic decidualization markers. Using qPCR we observed a reduction in *Bmp2* (P = 0.052) and *Wnt4* (P < 0.05) transcripts at GD5.5 in *Pgr^cre/+^; Ptgs2^f/f^* deciduae compared to controls (**Fig. 6A**). We also tested the decidual response of pseudo-pregnant control and mutant mice to an oil stimulus. We observed in comparison to the control uteri intraluminal oil stimulation of the *Pgr^cre/+^; Ptgs2^f/f^* uteri at GD2 1800h completely failed to elicit a decidualization response at GD5.5 (**Fig. 6B**). Taken together, our data suggest that stromal PTGS2 is crucial for post-implantation chamber growth, vessel remodeling surrounding the implantation chamber, and the initiation of decidualization, all of which are critical processes for successful pregnancy.

**Figure 6.**
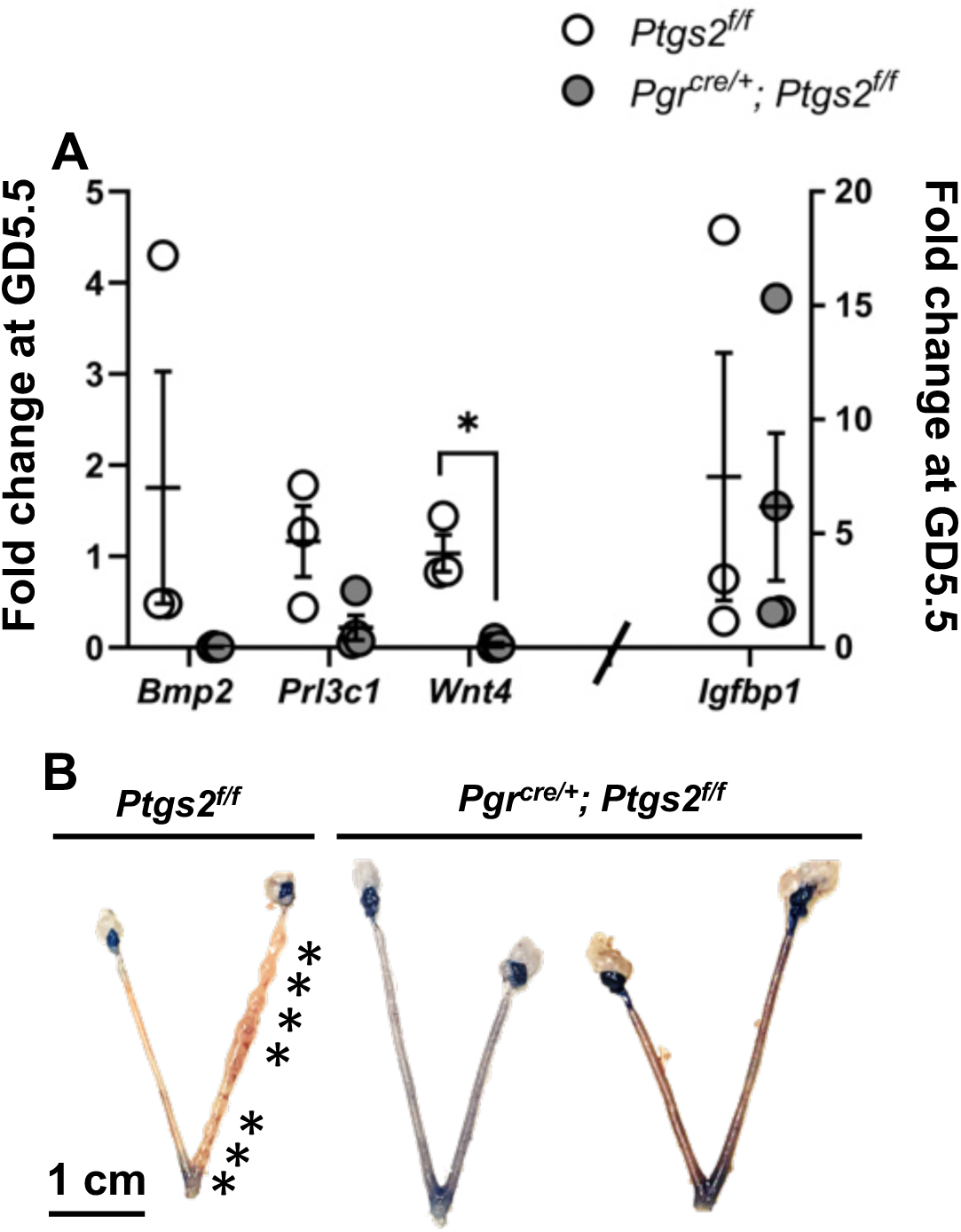
Decidualization failure in stromal deletion model of PTGS2 at GD5.5. Expression of decidualization markers measured by qRT-PCR at GD5.5 (A). Artificial decidualization induced by oil-stimulation for pseudopregnant mice at GD2 1800h and analyzed at GD5.5 (B). At least 3 mice for each condition were analyzed. Each dot represents one mouse. Median values shown. Data analyzed using unpaired parametric t-test. * P < 0.05. Scale bar, B: 1cm. Black asterisks: decidual sites.

## DISCUSSION

PTGS2-derived prostaglandins are functionally implicated in reproductive processes, including ovulation, fertilization, embryo implantation, and decidualization (Lim et al., 1997, Lim et al., 1999, Matsumoto et al., 2001, Kennedy, 1977). Despite these studies, there is still a debate in the literature regarding the role of PTGS2 in embryo implantation (Cheng and Stewart, 2003). In this study, we used different tissue-specific ablation models of PTGS2 and show that PTGS2 deletion in the uterine epithelium and endothelium does not impact pregnancy success. In contrast, deleting PTGS2 from the stroma results in post-implantation embryonic growth restriction, defective implantation chamber growth, and mid-gestation resorption. Our results highlight a role for uterine stromal PTGS2 in post-implantation stages of embryo development and initiation of decidualization but no critical role for PTGS2 in pre-implantation processes. During the drafting of this manuscript (Aikawa et al., 2024) published their observations using *Pgr^cre/+^; Ptgs2^f/f^* mice and their results are consistent with ours suggesting a role for stromal PTGS2 at the maternal-fetal interface. Given the debate on the role of PTGS2 function in murine pregnancy, consistent results with the tissue specific deletion highlight a role for PTGS2 function independent of mouse genetic background. The discussion below considers our study, as well as those by Aikawa et. al (Aikawa et al., 2024).

### Granulosa cell-specific deletion of PTGS2 does not produce ovulation and fertilization defects

PTGS2 is active in the ovaries during follicular development (Liu et al., 1997, Park et al., 2020), suggesting its importance during ovulatory processes. Clinical observations have reported luteinized unruptured follicle syndrome, characterized by the failure of follicle wall rupture despite a normal ovulatory cycle, in women who consume non-steroidal anti-inflammatory drugs such as indomethacin or selective PTGS2 inhibitors (Micu et al., 2011). This condition results in infertility(Qublan et al., 2006). In rodents, indomethacin treatment during proestrus disrupts the follicle rupture process, resulting in ovulation failure(Gaytán et al., 2002). Furthermore, both in vitro and in vivo studies have demonstrated that PTGS2 inhibition through indomethacin and NS-398 treatment inhibited LH hormone induction of PGE2 production and thus decreased ovulation rates in rats (Mikuni et al., 1998). *Ptgs2^-/-^* mice failed to produce PGs in response to gonadotropin stimulation and could not ovulate due to compromised cumulus expansion (Davis et al., 1999). This phenotype of failed ovulation occurs irrespective of mouse genetic background. These diverse lines of studies underscore the indispensable role of PTGS2 in ovulation. PGR and PTGS2 are co-expressed in the mural granulosa cells of the pre-ovulatory follicle following hCG stimulation (Zhang et al., 2023) and LH stimulation (Park et al., 2020). However, despite PTGS2 deletion in granulosa cells of the pre-ovulatory follicle and the corpus luteum of the ovary (Soyal et al., 2005), *Pgr^cre/+^; Ptgs2^f/f^* mice did not exhibit any ovulation failure. It is possible that *Pgr^cre^*may fail to delete *Ptgs2* in all granulosa cells, resulting in residual PTGS2 expression and function during ovulation. Alternatively, serum PGs synthesized outside the ovary, oviduct, and uterus may be responsible for the pro-inflammatory response resulting in ovulation. This will be a subject of future investigations.

### Uterine epithelial PTGS2 does not contribute to embryo spacing and on-time embryo implantation

The endometrial epithelium has been recognized as a source of the inducible PTGS2 and associated PGs, especially in the context of menstruation (Lundström et al., 1979). In addition, the epithelial and endothelial PGs are thought to regulate smooth muscle contraction and relaxation (Ruan et al., 2011, Félétou et al., 2011). Inhibiting PG synthesis results in embryo crowding in pregnant rats (Kennedy, 1977) and PGs are also critical for parturition (Reese et al., 2000, Aiken, 1972), highlighting a possible link between epithelial PTGS2 and muscle contractility for embryo spacing and parturition. Our expression studies did not detect epithelial or endothelial PTGS2 during the pre-implantation stage, although we did observe that PTGS2 is expressed in the luminal epithelium shortly after intraluminal stimulation with oil (Lim et al., 1997) and in the glands at the implantation chamber at GD5.5. Despite this, epithelial-only and epithelial and endothelial deletion of PTGS2 did not affect embryo spacing or on-time embryo implantation. Further, deletion of PTGS2 in the circular muscle in the *Pgr^cre/+^; Ptgs2^f/f^*did not affect embryo spacing, supporting that PTGS2 synthesized in the circular muscle, epithelium or endothelium is dispensable for uterine contractility critical for the initial phases of embryo movement.

### Uterine stromal PTGS2 is critical for decidualization success

Previous literature suggests that implantation and decidualization failure in *Ptgs2^-/-^* are not related to disruption in ovarian steroid levels or genes related to implantation success, such as *Leukemia inhibitory factor* (*Lif)* (Lim et al., 1997). Although progesterone levels were normal, we observed a significant reduction in *Lif* mRNA levels in our *Pgr^cre/+^; Ptgs2^f/f^* model. Reduced levels of *Lif* can explain the delay in implantation and may also contribute to the absence of decidualization response with an oil stimulus in this mutant. Delayed implantation may also explain the deviation of the embryo’s morphology compared to an elongated blastocyst at GD4. However, the absence of stromal PTGS2 at the anti-mesometrial pole of the implantation chamber in the *Pgr^cre/+^; Ptgs2^f/f^* model is the most likely cause of poor elongation of the implantation chamber and degradation of the embryos at GD5.5. A defective chamber likely results in a ripple effect of decreased vascular remodeling in the decidua surrounding the implantation chamber and reduction in the amount of decidualized stroma, leading to growth arrest in the embryo and failure of pregnancy progression. Our results suggest that elongation of the implantation chamber is critical for the transition of the embryo from an elongated blastocyst to an epiblast stage, highlighting a critical role for stromal PTGS2 in embryo-uterine communication at this stage of pregnancy.

Our results also highlight that once chamber formation begins and decidualization is initiated, the embryo is no longer needed for continuous expansion of the decidua. Even though 85% of embryos displayed severe growth retardation at GD5.5, decidual expansion continued until beyond GD8.5, and resorptions were only observed at a significant level at GD12.5 when extraembryonic tissue contributions are required for the formation of the placenta. These data are in line with other models of decidualization where oil and beads (Chen et al., 2011, Herington et al., 2009) can stimulate the initiation of decidualization, and the decidua continues to expand in the absence of embryonic contributions until mid-gestation. It has been proposed that decidualization with a bead or oil is different from embryo-induced decidualization (Herington et al., 2009). The *Pgr^cre/+^; Ptgs2^f/f^* mouse may be a good model to compare the growth of the decidua with and without a growing epiblast to explore the similarities and differences between the two decidualization processes.

Our data also highlights that even with complete ablation of stromal PTGS2 ∼50% of the embryos in the *Pgr^cre/+^; Ptgs2^f/f^* uteri continue to develop beyond mid-gestation and are also born. PTGS2 may permit implantation chamber growth beyond a certain length. If the chamber is stochastically able to grow beyond this length (due to PTGS1 upregulation or other factors such as the expanding decidua), then PTGS2 in the stroma may no longer be required. It is also possible that the embryos that display a delay in implantation are susceptible to the absence of stromal PTGS2 during the elongation of the chamber. However, these different hypotheses need to be tested to determine why some embryos continue to grow despite the absence of stromal PTGS2.

### Overlapping roles for PTGS1 and PTGS2 in murine implantation success and the role of mouse genetic background

*Ptgs1^-/-^* mice on a 129/B6 mouse background have 32% lower vascular permeability and significantly lower PG levels (specifically 6-keto-PGF1α and PGE_2_). These mice also display an upregulation of PTGS2 expression during the pre-implantation stage (Reese et al., 1999). This indicates that PTGS2 can compensate for the function of PTGS1 (Reese et al., 2000). When *Ptgs2* is inserted into the *Ptgs1* locus, PTGS2 can compensate for PTGS1 loss and rescue the parturition defect observed in *Ptgs1^-/-^*mice (Li et al., 2018b). However, on a C57Bl6 mouse background, when *Ptgs1* was placed in the *Ptgs2* locus, PTGS1 failed to compensate for PTGS2 function resulting in mice with implantation phenotypes similar to the *Ptgs2^-/-^* mice (Li et al., 2018b, Lim et al., 1997). It has been previously reported that on a mixed mouse genetic background, PTGS1 is upregulated in the *Ptgs2^-/-^* mice, and these mice exhibit improved fertility compared to *Ptgs2^-/-^* mice on a pure C57Bl6 mouse background (Wang et al., 2004a). In our studies (on a C57Bl6 background) and those by Aikawa et al (Aikawa et al., 2024) (mouse background not specified), the *Pgr^cre/+^; Ptgs2^f/f^* mice show ∼50% number of pups at birth. Consistency amongst our studies suggest a post-implantation role for PTGS2 independent of mouse genetic background. Aikawa et. al also showed that depletion of both PTGS1 and PTGS2 (*Ptgs1^-/-^*;*Pgr^cre/+^; Ptgs2^f/f^* mice, mouse background unknown), results in a complete failure of embryo implantation with embryos floating in the uterus (Aikawa et al., 2024). Since embryos were presumably normal in these mice, the cause for a complete absence of implantation could be lack of *Lif*, however this needs to be tested. All of these studies highlight the interconnected roles of PTGS enzymes and suggest that both PTGS1 and PTGS2 are critical for processes such as implantation and decidualization.

## Conclusions

Our study highlights that PTGS2-derived PGs necessary for implantation do not come from uterine epithelial and endothelial sources. Our work provides definitive proof that stromal PTGS2 at the base of the embryo implantation chamber is critical for both the growth of the embryo and the elongation of the implantation chamber. Further work is needed to understand how stromal PTGS2 depletion affects the decidualization response and vascular remodeling and why a certain percentage of embryos can escape this requirement and go through gestation. Overall, this study distinguishes between the pre-implantation and post-implantation roles of PTGS2 and provides a valuable model for investigating the role of stromal PTGS2 without the need for embryo transfer to study the initiation of the decidualization process and how it relates to pregnancy success.

## METHODS

### Animals

We generated the *Ptgs2* conditional deletion mice by breeding C57/bl6 *Ptgs2^f/f^* (Ishikawa and Herschman, 2006) with C57/bl6 *Ltf^cre/+^* (Daikoku et al., 2014), mixed genetic background *Pax2^cre/+^*(Ohyama and Groves, 2004), or C57/bl6 *Pgr^cre/+^* (Soyal et al., 2005) mice (**Table 1**). For pregnancy studies, we set adult females at 6-10 weeks to mate with fertile males. For *Ltf^cre/+^; Ptgs2^f/f^*, we mated them between 10-12 weeks, as PTGS2 deletion occurs in the adult females (Daikoku et al., 2014). To create pseudopregnancy, we mated females with vasectomized males. The appearance of a vaginal plug was identified as a gestational day (GD) GD0 1200h. We euthanized mice at several stages, including GD3 1200h and GD3 1800h, GD4 1800h, GD5.5, GD8.5, and GD12.5, or mice were allowed to go to term. We performed GD5.5, GD8.5, and GD12.5 dissections between 1300h and 1500h on the dissection day. To induce an artificial decidualization, we used a non-surgical embryo transfer (NSET) device, where we transferred 1 µl sesame oil and 3 µl PBS to a pseudo-pregnant mouse on either GD2 1800h or GD3 0800h. We euthanized the oil-stimulated pseudo-pregnant mice at GD3 1200h or GD5.5. For GD4 and GD5, we euthanized the animals 10 minutes after 0.15 ml intravenous injection of 1.5% of Evans blue dye (MP Biomedicals, ICN15110805). All mice were maintained on a 12-hour light/dark cycle, and all mouse studies and protocols were approved by the Institutional Animal Care and Use Committee at Michigan State University.

### Whole-mount immunofluorescence staining

As described previously (Arora et al., 2016, Flores et al., 2020, Madhavan et al., 2022) for whole-mount staining, we fixed dissected uteri in a mixture of cold DMSO: Methanol (1:4). We hydrated the samples in a (1:1) methanol: PBST (PBS, 1% triton) solution for 15 minutes, followed by a 15 minutes wash in PBST. We then placed the samples in a blocking solution (PBS, 1% triton, and 2% powdered milk) for 1 hour at room temperature followed by incubation with primary antibodies (**Supplementary Table 1**) in the blocking solution for seven nights at 4°C. After washing with 100% PBST solution for 2X15 minutes and 4X45 minutes, we incubated the samples with Alexa Flour-conjugated secondary antibodies for three nights at 4°C (**Supplementary Table 1**). Following the incubation, we washed the samples with PBST for 2X15 minutes and 4X45 minutes and incubated the samples at 4°C overnight with 3% H2O2 diluted in methanol. Finally, we washed the samples with 100% methanol for 3X30 minutes and cleared the tissues overnight with benzyl alcohol: benzyl benzoate (1:2) (Sigma-Aldrich, 108006, B6630).

### Cryo-embedding, cryo-sectioning, and immunostaining

As described previously (Granger et al., 2023) we fixed uterine tissues in 4% PFA (paraformaldehyde) for 20 minutes and then incubated the samples with fresh 4% PFA overnight at 4^°^C. The tissues were then washed with PBS for 3X5 minutes and then incubated in 10% sucrose prepared in PBS at 4°C overnight. We then transferred the samples to 20% and 30% sucrose solutions in PBS for 2-3 hours each at 4°C. Then we embedded the samples in tissue-Tek OCT (Andwin Scientific, 45831) and stored them at -80°C. Cryo-sections of 7µm thickness were mounted on glass slides (Fisher, 1255015). For the immunofluorescent staining, we allowed the slides to air dry for 15 minutes and then washed them with PBS for 3X5 minutes, and blocked with PBS + 2% powdered milk + 1% triton solution for 20 minutes. After additional PBS for 3X5 minutes washes, we stained the slides with primary antibodies (**Supplementary Table 1**) and incubated them at 4°C overnight. The next day we washed the slides with PBS for 3X5 minutes and incubated them with secondary antibodies and Hoechst (**Supplementary Table 1**) for 1 hour at room temperature. Finally, after PBS washes, we added 2 drops of 20% glycerol in PBS to the slides followed by sealing the sections with glass coverslips.

### In situ hybridization

We performed in situ hybridization on uterine sections using the RNAscope 2.5 HD Assay-RED kit (ACD Bio, 322350), which also has immunofluorescence capabilities, as described previously (Granger et al., 2023). We aimed to detect *Lif* mRNA associated with the uterine glands at GD3 1800h. To detect *Lif*, we used the Mm-Lif probe (ACD Bio, 475841), and to label uterine glands, we included immunostaining for FOXA2 (**Supplementary Table 1**). The entire 3-day protocol was carried out according to the protocols provided by ACD Bio (322360-USM, MK 51-149 TN).

### Serum progesterone measurement

After euthanizing the mouse, we collected 200-500 µl of blood samples and left them at room temperature for 30 minutes. Then, we centrifuged the samples for 15 minutes at 2000 g, carefully separated the supernatant, and immediately saved the samples at -20°C. Following sample collection and preservation, we sent the samples to a Ligand Assay and Analysis Core Laboratory in Charlottesville, VA, to determine progesterone levels. Samples were diluted at a ratio of 1:4, tested in triplicate to ensure accuracy, and the results were reported in ng/ml.

### Oviduct flush and in vitro embryo culture

For oviduct flush at GD1 1200h, we euthanized the female mice, excised both oviducts and placed them in warm (37°C) M2 medium (Sigma-Aldrich, M7167). We flushed each oviduct with approximately 300 – 500 ul of pre-warmed (37°) M2 medium using a blunted 30-gauge needle attached to a 1ml syringe. We collected embryos and unfertilized eggs using a mouth pipette with a pulled glass capillary. After washing them 2 to 3 times in warm (37°C) KSOM medium (Cyrospring), we incubated them in 400-600 µl drop of KSOM media and placed them in a 37°C jacketed incubator. We monitored embryonic development daily for 72 hours and recorded the number of embryos reaching 4-cell, 8-cell, morula, and blastocyst stages (Frum and Ralston, 2019).

### RNA Isolation, cDNA Synthesis, and quantitative PCR

We isolated uterine decidual tissues at GD5.5, snap-froze, and stored the samples at -80 °C. We isolated total RNA from tissues using the Trizol reagent (Invitrogen, 15596019). Briefly, we homogenized the tissues in 1 ml TRIzol solution using the Bead Mill 4 homogenizer (Thermo Fisher Scientific). Following phase separation with 500 µl chloroform, RNA was precipitated with isopropanol and washed with 75% ethanol. Then, we suspended the RNA in 50-100 µl RNase-free water (Invitrogen, AM9922). We measured the RNA concentration and purity using a NanoDrop 2000 spectrophotometer (Mettler Toledo) with a concentration of at least 250 ng/µl. We performed first-strand cDNA synthesis from 1 ug RNA using reverse transcriptase enzyme (Promega, PRA5003). For qRT-PCR, we designed the primers using the primer3Plus and NCBI website (**Supplementary Table 2**). We carried the qRT-PCR reactions in triplicate for each sample using the Quantstudio 5 Real-Time PCR system (Applied Biosystems) with a total reaction volume of 20 µl (10 µl SYBER Green (Thermo Fisher Scientific, A25742), 7.4 µl Rnase and Dnase free water, 1.6 µl primer, and 1 µl cDNA). We used the comparative CT (ΔΔCt) method for gene expression analysis. We calculated the ΔCt for each sample by subtracting the Ct value of the *Rpl19* gene from the Ct value of the target gene. We calculated the ΔΔCt by subtracting the mean ΔCt of the control group from the ΔCt of each sample. Fold change was calculated as 2^(-ΔΔCt) (Livak and Schmittgen, 2001).

### Confocal Microscopy

We used a Leica SP8 TCS white light laser confocal microscope utilizing 10x air to image whole uterine tissues or 20X water objective and a 7.0 um Z stack or system-optimized Z stack to image the samples (Madhavan et al., 2022). Upon imaging, we imported the files (.LIF format) into Imaris v9.2.1 (Bitplane; Oxford Instruments, Abingdon, UK) 3D surpass mode. We created 3D renderings using surface modules.

### Image Analysis

#### Implantation chamber, luminal epithelium, and embryo visualization

To visualize the implantation chamber, we used the CDH1 fluorescent signal for the luminal epithelium surface and the FOXA2 fluorescent signal for uterine glands. We isolated the luminal epithelium by subtracting the FOXA2-specific signal from the CDH1 signal. We used the Hoechst signal to locate embryos based on the inner cell mass (ICM) signal, and we used the 3D rendering surface in IMARIS software to create the embryo surfaces. We used the measurement function in Imaris to measure the length of the implantation chamber.

#### *Lif* quantitation

As described (Granger et al., 2023), we used the FOXA2 signal to generate 3D surfaces of the glands’ nuclei via the 3D surface function within the IMARIS software. Subsequently, we used the IMARIS masking function to produce a distinct channel for the *Lif* signal that lies beneath the previously established uterine gland 3D surface. Based on the new channel for the *Lif* signal, we created a new 3D surface of *Lif*. Following the creation of the 3D surfaces, we used the statistics function of Imaris to determine the 3D surface volume of both the glands and *Lif*. We used Microsoft Excel to calculate the *Lif* volume per uterine gland volume and plotted the results as *Lif* volume per uterine gland volume (FOXA2 signal) with normalized units.

#### Vessel density around the anti-mesometrial pole of the implantation chamber

We created a 3D rendering surface of blood vessels using a CD31 fluorescent signal and generated a channel in Imaris software to mask the surface of the blood vessels. For image segmentation, we imported 14 µm of the masked channel of vessels to ImageJ after adjusting the scale and applying the threshold function. Using vessel analysis and Mexican Hat Filter Plugins in ImageJ (https://imagej.net/), we calculated the density and diameter of the blood vessels in the embryo implantation and inter-implantation site. For vessel density the data is reported as the percentage of area occupied by blood vessels.

### Statistical Analysis

We used Graph Pad Prism (Dotmatics; GraphPad, La Jolla, CA, USA) and Microsoft Excel to analyze the statistical differences between the treatment groups and plot our graphs. To analyze the difference between the two treatment groups, we employed the unpaired parametric two-tail t-test. First, we tested the data for homogeneity of the variance between the two treatments. If the variances were equal, we proceeded with a parametric two-tailed t-test. If the variances differed, we used the Mann-Whitney U-test to compare the two treatment groups. We considered the data statistically different for P value < 0.05 or less.

## ACKNOWLEDGEMENTS

We thank Dr. Harvey R. Herschman and Dr. Srinivasa Reddy at UCLA for providing *Ptgs2*-floxed mice. We thank Dr. Gregory Burns, Dr. Nataki Douglas, Dr. Shuo Xiao, and Dr. Asgerally Fazleabas for critical discussions related to the project.

## AUTHOR CONTRIBUTIONS

NM and RA conceptualized the study and designed the experiments. NM executed experiments. NM and RA validated the data and performed the analyses. NM and RA wrote and edited the manuscript. All authors reviewed and accepted the final version of the manuscript.

---

**Supplementary Figure 1:**
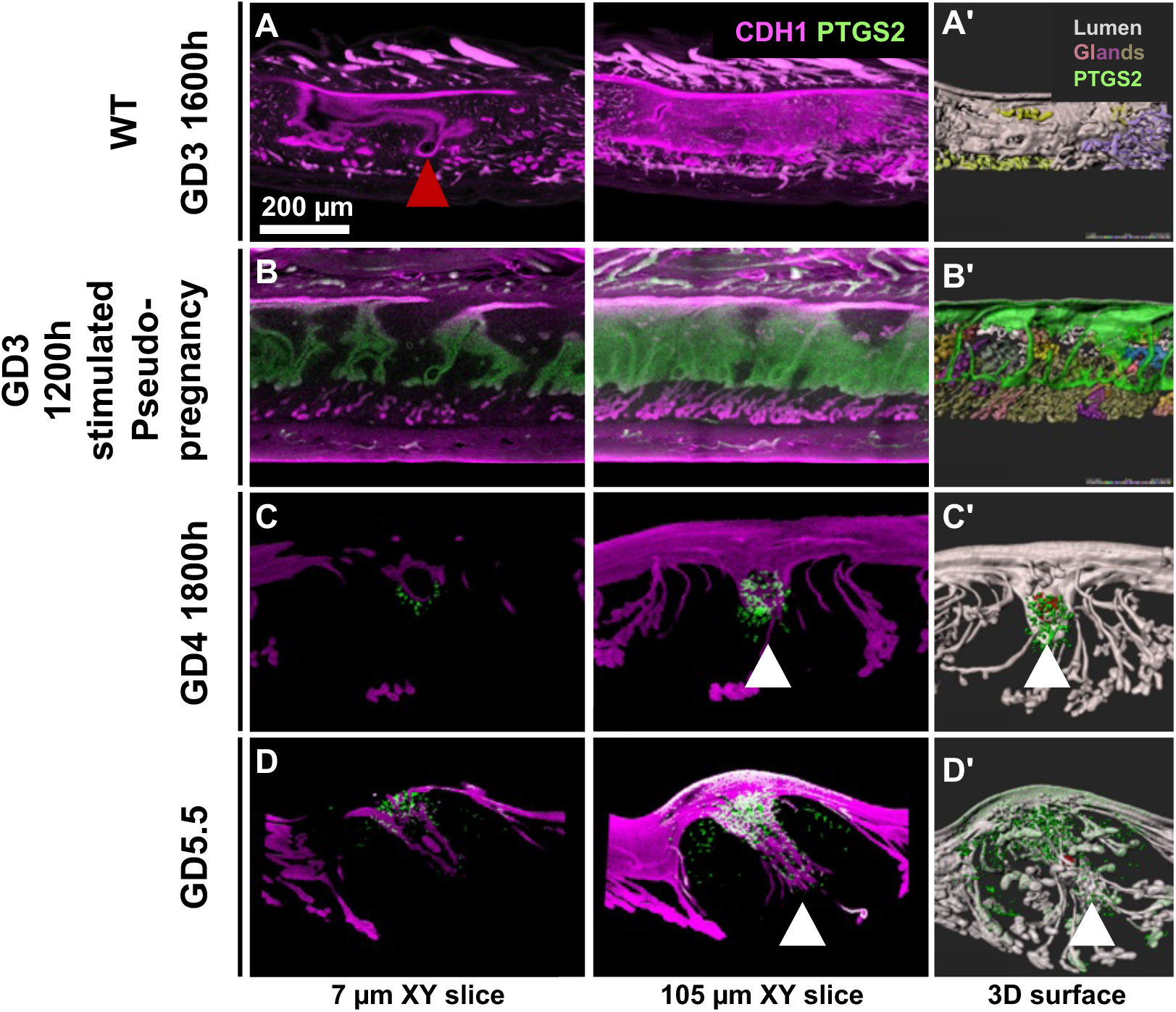
Timeline of uterine PTGS2 expression during peri-implantation stages and in an oil-stimulated pseudopregnancy. CDH1 and PTGS2 expression in pregnant wild-type uteri at GD3 1600h (A). PTGS2 expression in CDH1 positive cells in oil-stimulated pseudo pregnant uteri at GD3 1200h, 4h after oil stimulation (B). 3 different regions from at least 8 uterine horns were evaluated. PTGS2 expression in the subepithelial stroma surrounding the embryo implantation chamber at GD4 1800h (C). PTGS2 expression in the mesometrium pole and the uterine glands of the embryo implantation chambers at GD5.5 (D). At least 2 implantation sites from at least 3 uterine horns were analyzed. 7µm XY slice (A, B, C, D). 105 µm XY slice (A’, B’, C’, D’). Scale bar, A-D’: 200 µm. GD: gestational day; Red arrowhead: embryo; White arrowheads: embryo implantation chamber.

**Supplementary Figure 2.**
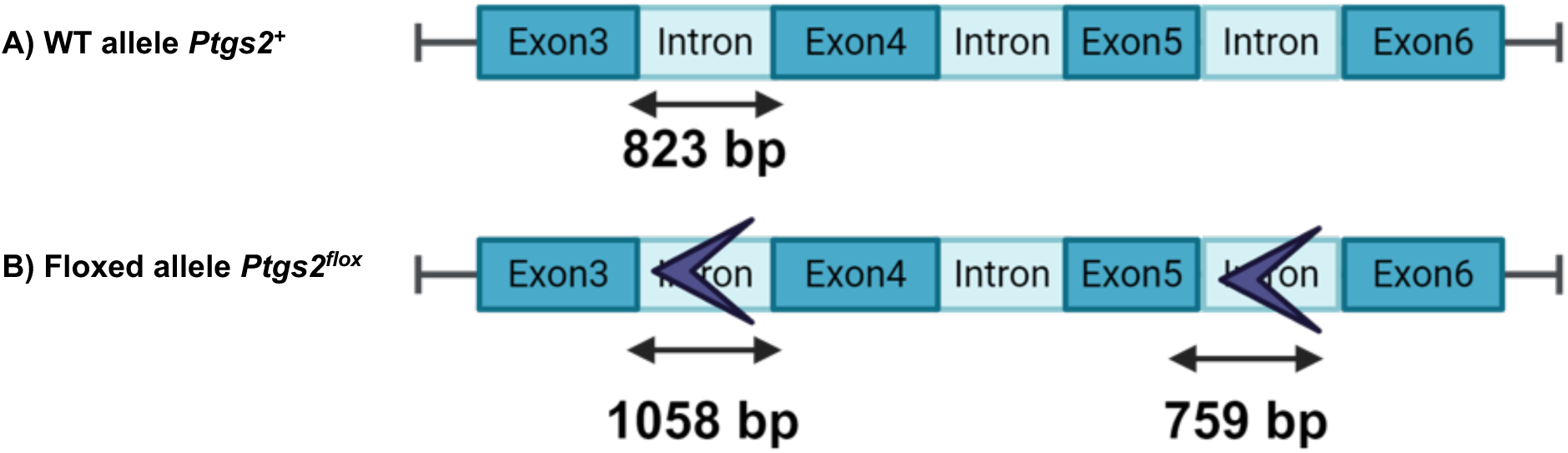
Utilizing the cre-lox recombinase system to specifically delete PTGS2 from the murine reproductive tract. The diagram displays the wild-type allele of *Ptgs2* (A) and the floxed allele of *Ptgs2* with the lox-p sites inserted in introns 3 and 5 (B).

**Supplementary Figure 3:**
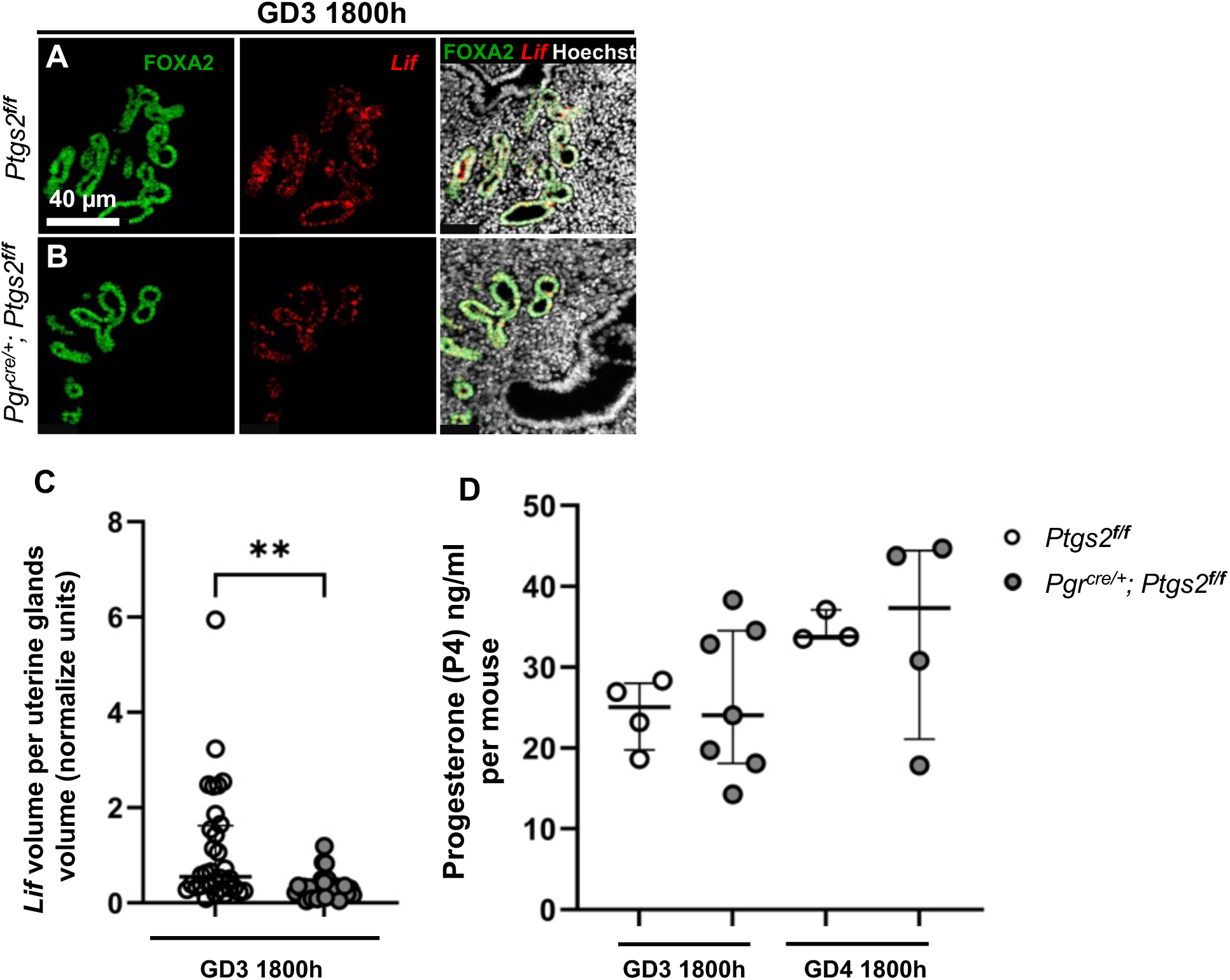
*Pgr^cre/+^; Ptgs2^f/f^* uteri display a reduction in preimplantation *Leukemia Inhibitory Factor*. *Leukemia inhibitory factor* (*Lif) expression in* FOXA2+ glandular epithelium cells in *Ptgs2^f/f^* and *Pgr^cre/+^; Ptgs2^f/^*at GD3 1800h (A, B). Quantification of *Lif* volume normalized to FOXA2+ glandular epithelium volume at GD3 1800h per uterine section in the two groups (C). At least 4 mice and 28 sections were analyzed for each group. Each dot represents one uterine section. Median values shown. Data analyzed using unpaired parametric t-test.** P < 0.01. Progesterone serum levels in *Ptgs2^f/f^* and *Pgr^cre/+^; Ptgs2^f/f^* at GD3 1800h and GD4 1800h (D). At least n=3 mice per genotype were analyzed. Each dot represents one mouse. Median values shown. Data analyzed using unpaired parametric t-test. No significant difference observed. Scale bar, A-B: 40 µm.

**Supplementary Table 1.** Primary and secondary antibodies used in the study.

| Primary antibody | Dilution | Stained tissues |
| --- | --- | --- |
| Rat-anti-CDH1 (M108, 108006, B6630) | 1:500 | Luminal epithelium |
| Rabbit anti-FOXA2 (Abcam, ab108422) | 1:500 (WM)<br>1:200 (sections) | Glandular epithelium |
| Rabbit anti-PTGS2 (Abcam, ab16701) | 1:500 | PTGS2-positive cells |
| Armenian-hamster anti-CD31 (DSHB, AB_2161039) | 1:200 | Endothelial cells |
| Secondary antibody | Dilution |  |
| Goat anti-rat 647 (A21247, Invitrogen, Carsbad, CA, USA) | 1:500 |  |
| Donkey anti-rabbit 555 (A31572, Invitrogen) | 1:500 |  |
| Donkey anti-rabbit 555 (A31572, Invitrogen) | 1:500 |  |
| Nuclear marker | Dilution |  |
| Hoechst (Sigma Aldrich, B2261) | 1:500 |  |

**Supplementary Table 2.**
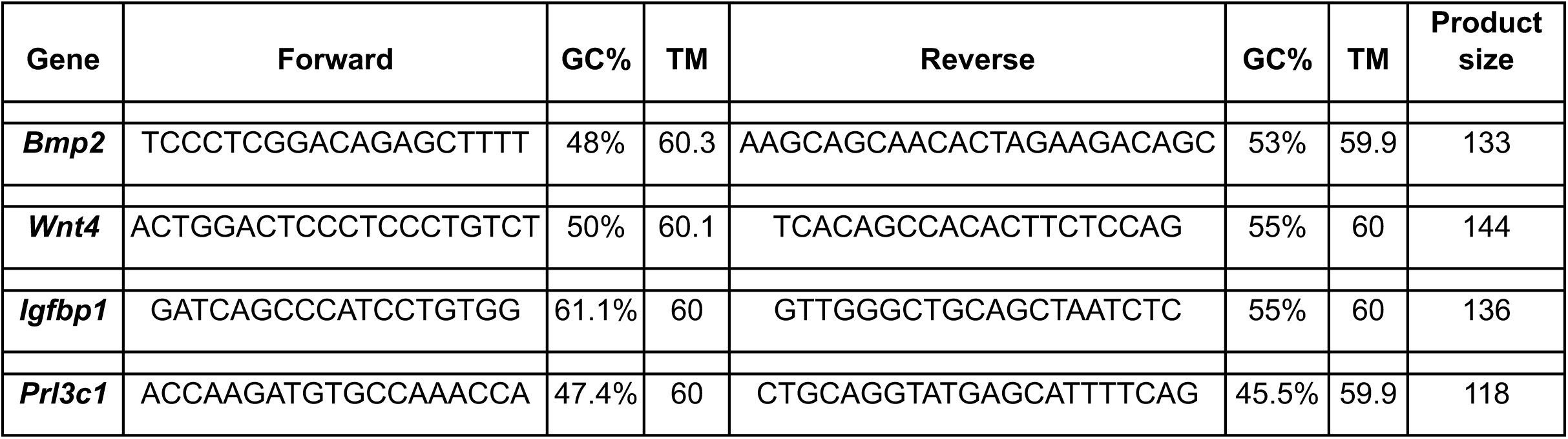
Primers sequences for Quantitative real-time polymerase chain reaction (PCR). Forward and reverse primer sequences for decidualization genes (*Bmp2*, *Wnt4*, *Igfbp1*, *Prl3c1*). TM: melting temperature.

